# Muscarinic M1 receptor modulation of synaptic plasticity in nucleus accumbens of wild-type and fragile X mice

**DOI:** 10.1101/270256

**Authors:** Daniela Neuhofer, Olivier Lassalle, Olivier J Manzoni

## Abstract

We investigated how metabotropic Acetylcholine receptors control excitatory synaptic plasticity in the mouse nucleus accumbens core. Pharmacological and genetic approaches revealed that M_1_ mAChRs trigger multiple and interacting forms of synaptic plasticity. As previously described in the dorsal striatum, moderate pharmacological activation of M_1_ mAChR potentiated postsynaptic NMDARs. The M_1_-potentiation of NMDAR masked a previously unknown coincident TRPV1-mediated long-term depression (LTD). In addition, strong pharmacological activation of M_1_ mAChR induced canonical retrograde LTD, mediated by presynaptic CB1R. In the *fmr1*-/y mouse model of Fragile X, we found that CB1R but not TRPV1 M_1_-LTD was impaired. Finally, pharmacological blockade of the degradation of anandamide and 2-arachidonylglycerol, the two principal eCBs restored *fmr1*-/y LTD to wild type levels. These findings shed new lights on the complex influence of Acetylcholine on excitatory synapses in the nucleus accumbens core and identify new substrates of the synaptic deficits of Fragile X.

## Introduction

Acetylcholine is a major neurotransmitter and modulator in the CNS, acting via ionotropic nicotinic and metabotropic muscarinic receptors it is involved in a plethora of cognitive and executive functions^1^.

Five distinct mAChR subtypes (M_1_–M_5_), are expressed in the brain^2,3^ and quantitative autoradiographic studies have demonstrated that the striatum has one of the highest concentrations of muscarinic receptors^4^ highlighting the importance of muscarinic signaling in the basal ganglia. The role of dorso-striatal cholinergic transmission in the control of voluntary movement is well established^5^. The ventral part of the striatum, the nucleus accumbens has been conceptualized as the « gatekeeper » of the basal ganglia, because it is ideally positioned to integrate signals originating from limbic and cortical areas and modulate reward-related motor output^6^. The accumbens has been extensively studied in the context of drug abuse and addiction related behaviors^7^. More recently, its role in rewarding social behaviors and social interactions has been highlighted^8–10^. Muscarinic and nicotinic receptors in the accumbens are necessary for the acquisition of appetitive tasks^11^, food and drug satiety^12^. How cholinergic inputs modulate glutamatergic synaptic transmission onto medium spiny neurons (MSN) remains poorly understood.

M_1_ mAChR activation triggers long-term depression (LTD) in the perirhinal cortex^13^, the visual cortex ^14^, the hippocampus^15,16^, the prefrontal cortex^17^ as well as axonal signal processing^18^. In contrast to the dorsal striatum, how mAChR modulate synaptic plasticity in the accumbens remains largely unknown.

Cholinergic dysfunction has been implicated in the pathophysiology of schizophrenia, mood disorders as well as neurodegenerative disorders including Alzheimer and Parkinson diseases^1,19,20^. Fewer studies have addressed the implication of the cholinergic system in Fragile X syndrome (FXS), the most common monogenetic cause of inherited intellectual disability and a leading cause of autism^21–23^. The disease is caused by mutation of a single X-linked gene called *fmr1*^24^. The Fragile X mental retardation protein (FMRP) is a 71 kDa protein which regulates the transport and translation of more than 850 mRNAs in the brain and especially in synapses^25–27^. In humans with FXS, the loss of FMRP results in a variety of neurological symptoms widely associated with dysfunctional synaptic plasticity in critical brain regions such as the cortex, hippocampus, and amygdala^28, 29^. In the *fmr1*-/y mice model of FXS, structural and functional deficits have been reported in multiple brain areas, most notably the hippocampus, the cortex but also the striatum and accumbens^30–34^. Although, acetylcholine plays a key role in arousal and reward and FXS patients commonly show symptoms in associated behaviors^28, 35^, how acetylcholine-accumbens plasticity is affected in *fmr1-/y* mice is currently not known.

Here we used pharmacological methods to explore Acetylcholine-dependent synaptic plasticity and its underpinnings in the accumbens core region of wild-type and *fmr1-/y* mice. We report that two types of M_1_ mAChR-mediated LTD and one LTP cohabit at excitatory synapses onto accumbens core MSN. Moderate pharmacological activation of M_1_ mAChR induces both a TRPV1-mediated LTD and a potentiation of NMDAR, two phenomena that occlude mutually. In response to strong activation, M_1_ mAChRs induce a CB1R-mediated retrograde LTD. Finally, we show that CB1R-but not TRPV1-mediated M_1_-LTD was affected in *fmr1*-/y mice and that pharmacological blockade of the degradation of anandamide and 2-arachidonylglycerol, the two principal eCBs restored LTD in the Fragile X mouse model.

The results provide a previously unidentified link between M_1_ mAChR-mediated accumbal synaptic plasticity and cognitive dysfunction in Fragile X and suggest the cholinergic system as a novel therapeutic target.

## Results

### Direct activation of muscarinic M1 receptors induces LTD in the accumbens core

Acute cholinergic stimulation induces synaptic plasticity in several cortical areas^17,36^. Striatal medium spiny neurons (MSN) receive cholinergic innervation from the brainstem^37^ and local giant cholinergic interneurons^38^. We first tested the hypothesis that G-protein coupled muscarinic Ach receptors (mAChR) can modulate excitatory synapses in the accumbens.

Recording field EPSPs from MSN in the accumbens core, we first challenged slices from adult wild type mice with brief (10 minutes) applications of the large spectrum muscarinic agonist Carbachol. Figure 1A shows the individual field responses in two representative experiments. Bath perfusion with 10 μM Carbachol induced a short lasting and fully reversible depression (STD), which returned to baseline levels after 20 minutes. In contrast, bath-perfusion of 100 μM Carbachol induced a sustained long-term depression (LTD) of synaptic efficacy in the accumbens core. Figure 1B summarizes the average field responses for the three different concentrations tested. All three concentrations induced significant STD in comparison to baseline response (1μM: 79.27±5.373, p=0.006 n=8; 10μM:52.89±3.730, p<0.0001, n=9; 100μM: 42.22 ±3.249, p<0.0001 n=13; one-sample t-test). There was a concentration-dependent difference in the amount of STD (one-way ANOVA p < 0.0001 with Holm-Sidak’s multiple comparisons test: 1μM vs 10μM p=0.0002; 1μM vs 100μM p < 0.0001; 1μM vs 10μM; 10μM vs 100μM p=0.0216). From our results, it is clear that LTD was triggered solely in response to the highest dose of Carbachol (1μM: 96.48±6.677, p=0.6146; 10μM: 107.3±4.661, p=0.1555; 100μM: 76.90±4.190,p<0.0001; one-sample t-test). Figure 1C shows the lack of correlation between the magnitude of STD and LTD at all three concentrations of Carbachol tested (1μM p=0.8653; 10μM p=0.4967; 100μM p=0.2187; Pearson’s correlation).

**Figure 1:**
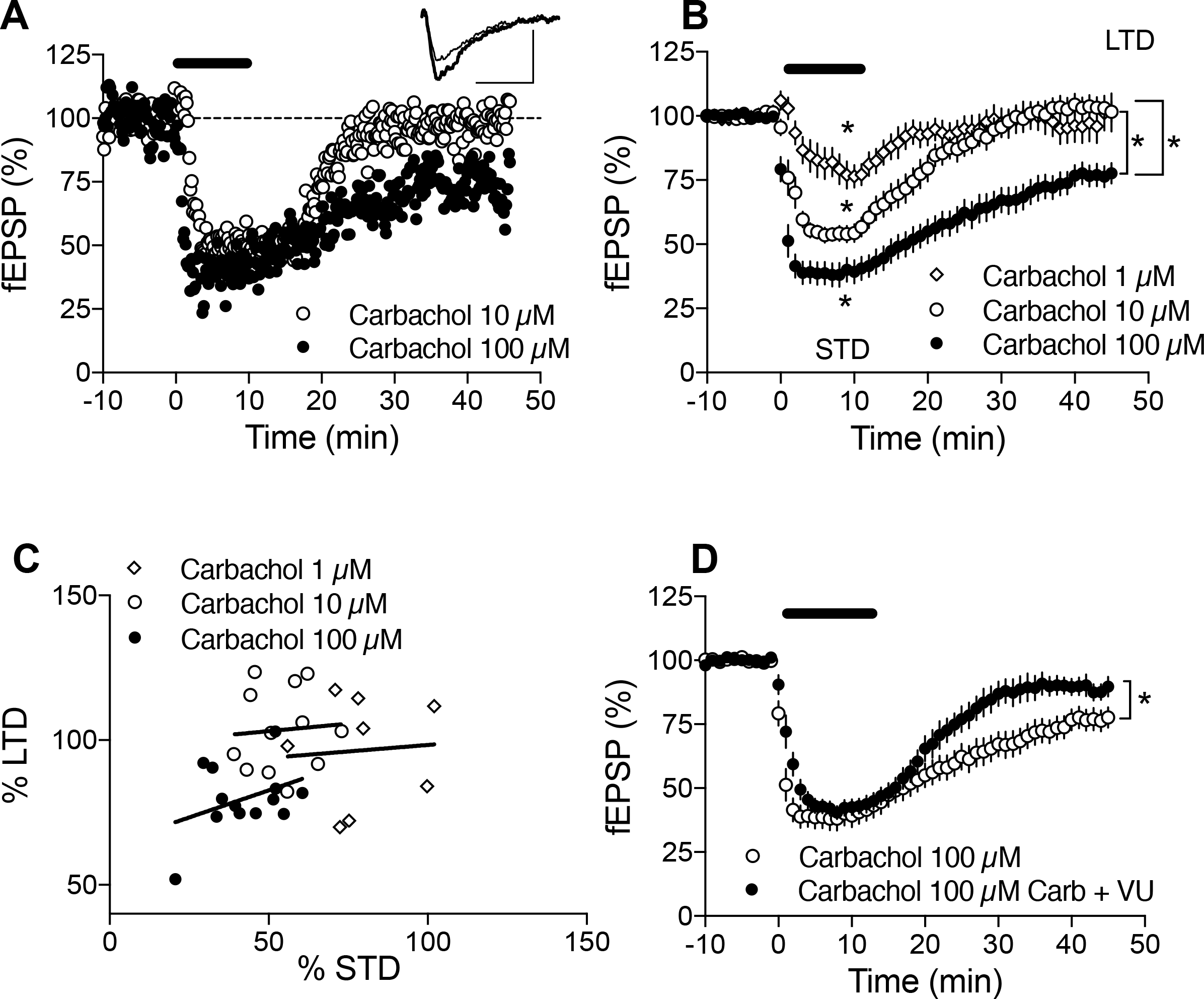
Direct pharmacological activation of M1 AChR triggers STD and LTD in the nucleus accumbens core. A) Representative field recording showing the effects of 10μM and 100μM Carbachol. The lowest dose (10μM) induced a strong but transient depression of synaptic responses (short-term depression, STD). The highest concentration of Carbachol (100μM) induced a robust LTD. B) Averaged fEPSPs for three different Carbachol concentrations (1mM, n=8; 10μM n=9; 100μM n=13). All three concentrations induced STD but only 100 μM Carbachol induced LTD. C) Pearson’s correlation showed no dependence of LTD magnitude on STD. D) 100 μM Carbachol mediated LTD was highly sensitive to the M1 Antagonist VU0225035. n=12 \**P* < 0.05.

In the medial and prelimbic prefrontal cortices, Carbachol induced LTD is mediated by M_1_ mAChR^17,36,39^. Thus, accumbens slices were incubated with the M_1_ specific receptor antagonist VU0225035 (10μM). Figure 1D shows that LTD was inhibited by the M_1_ mAChR antagonist (92.92±4.292, n=12, p=0.1270, one-sample t-test), showing that activation of M_1_ mAChR induces LTD in the accumbens core as demonstrated previously in the prefrontal cortex^17,36,39^.

### Potentiated synaptic NMDA currents mask “low Carbachol” LTD

In contrast with previous studies, including one from our laboratory demonstrating LTD induced by low concentrations of Carbachol in the prefrontal cortex (PFC)^17,39^, we were surprised to observe that bath application of 10 μM Carbachol was not sufficient to induce LTD. Such discrepancy could be due to low M_1_R expression, poor M_1_R -coupling efficiency to downstream effectors or result from multiple compensating/antagonizing M_1_R-mediated synaptic effects. Noteworthy, activation of M_1_R potentiates NMDAR currents and offsets LTD in the dorsal striatum^5,40^. To test if a similar process occurs in the accumbens core, we simply recorded NMDAR-mediated fEPSP in ACSF containing 0 Mg^++^ and 100 μM CNQX to block ionotropic glutamate receptors fast synaptic potentials (i.e. mediated by AMPAR / KAR). fEPSP recordings were chosen to allow for the direct comparison with our LTD experiments and their robustness to pharmacological treatments. Figure 2A shows the average NMDAR field recordings and the inset illustrates two averaged field responses (pre and post Carbachol application): 10μM carbachol induced a rapid short-term depression of NMDAR mediated fEPSPs (66.23±4.884, n=5, p=0.0023, one-sample t-test; Figure 2B) followed by a long-term potentiation (LTP) after drug washout (137±12.6, p=0.0425, one-sample t-test), whereas 100μM Carbachol also induced rapid short-term depression of NMDAR fEPSPS (43.58±10.67, n=8, p=0.2156, one-sample t-test) followed by a trend towards long-term depression (LTD) after drug washout (81.22±13.79, p=0.2156, one-sample t-test). Thus, NMDAR-mediated synaptic responses in the accumbens are potentiated by mAChR, as previously reported in the dorsal striatum^5,40^.

**Figure 2.**
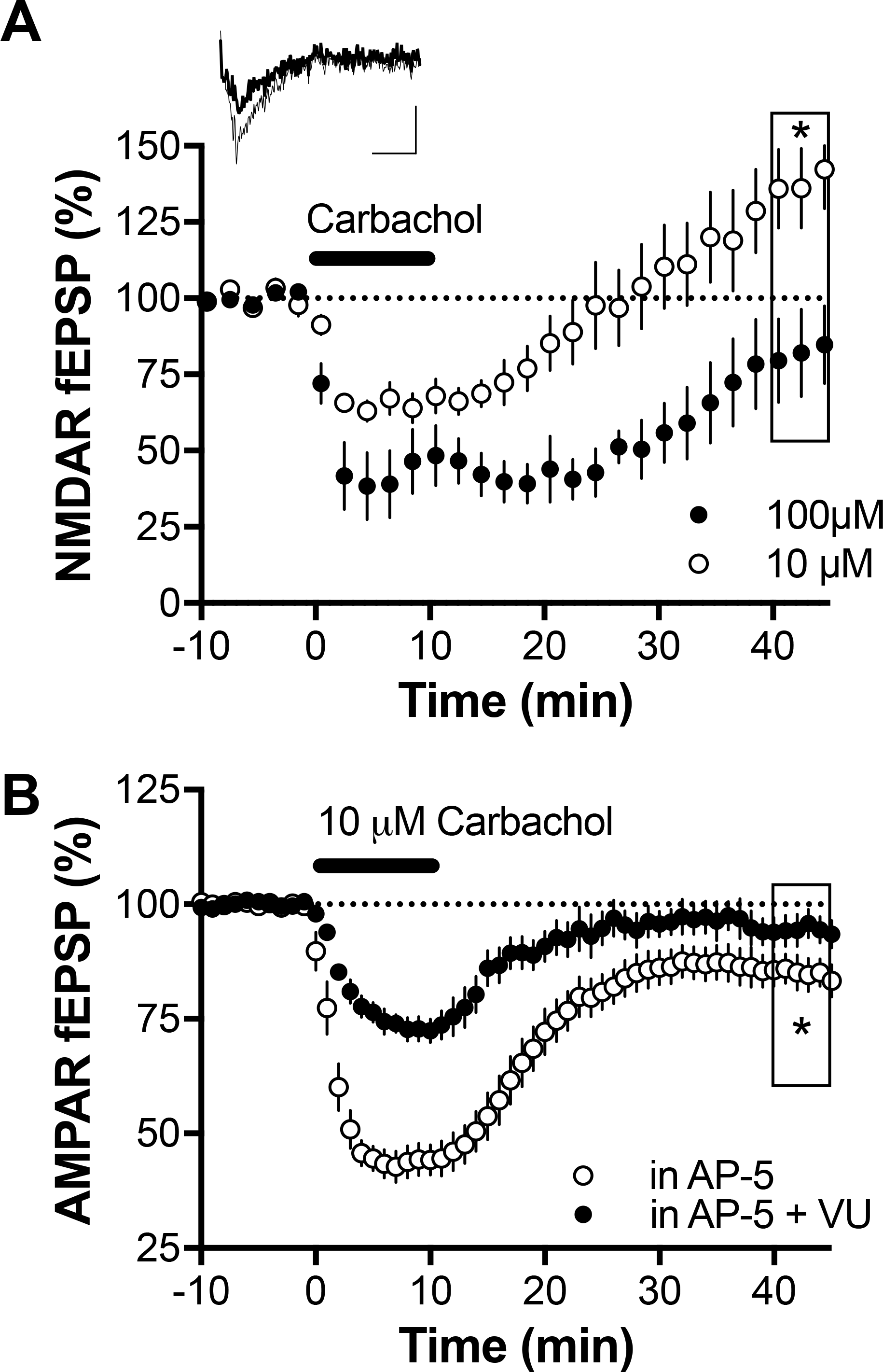
NMDAR modulate mAChR-induced synaptic plasticity. A) Averaged field recordings of NMDAR responses. In response to bath-application of 10μM Carbachol, the transient depression was followed by a marked LTP of NMDAR-fEPSP. In response to bath-application of 100μM Carbachol, the transient depression was followed by a trend towards LTD of NMDAR-fEPSP Inset: representative field response before and after 10μM Carbachol application. B) Average field recordings of AMPAR responses. 10 μM Carbachol mediated LTD was unmasked after blocking NMDAR with dAPV. This LTD was blocked with the Mi specific receptor antagonist VU0225035 (50μM). . * p<0.05

We hypothesized that LTP of NMDAR synaptic potentials might mask the LTD of AMPAR fEPSPs. We reasoned that NMDAR antagonism could unmask LTD in slices perfused with low Carbachol. In support of this scenario, bath perfusion of 10μM Carbachol in presence of the NMDAR antagonist D-AP5 (50μM) now induced a significant LTD (85.28±3.365, n=15, p=0.0006, one-sample t-test). This LTD was blocked in accumbens slices incubated with the M_1_ specific receptor antagonist VU0225035 (10μM) (94±3.103, n=8, p=0.1126, one-sample t-test, Figure 2B).

Together our data show that moderate activation of M_1_ mAChR with 10μM Carbachol induces concomitant AMPAR LTD and NMDAR LTP. Although “low Carbachol” largely modulates AMPAR and NMDAR function, the change in synaptic transmission can only be unmasked by blocking NMDAR.

### TRPV1 receptors -not CB1R - mediate “low Carbachol” LTD

Different LTD pathways allow a single neuron to engage either presynaptic CB1R or postsynaptic TRPV1 receptors^41^. It has been long established that M_1_ mAChR can engage the production of endocannabinoids (eCB) to consequently modulate short and long term synaptic plasticity^42,43^. In the bed nucleus stria terminalis^41^ and also the accumbens, eCB engage presynaptic CB1R and /or postsynaptic TRPV1R depending on cell type and stimulation patterns^44,45^.

Hence, we explored the locus of LTD expression and the mechanism of the low Carbachol LTD. A series of experiments was performed in the presence of D-AP5 to block NMDAR and unmask LTD (Figure 3). We found that induction of low Carbachol LTD was abolished in slices incubated with the selective TRPV1 receptor antagonist AMG9810 (98.21±5.984, n=5, p=0.7801, student’s t-test; Figure 3A-C). However, low Carbachol LTD was unaffected by the CB1R antagonist SR141716A (83.64±5.932, n=10, p=0.0220, student’s t-test; Figure 3C). We verified the locus of LTD by simply quantifying the changes in the paired-pulse ratio from the field excitatory responses (Figure 3A, lower trace). The paired-pulse ratio quickly returned to baseline (p=0.5818, student’s t-test), pointing toward a postsynaptic expression mechanism of LTD expression. Taken together these data strongly suggest that low Carbachol induces a postsynaptic LTD mediated by TRPV1 receptors.

**Figure 3:**
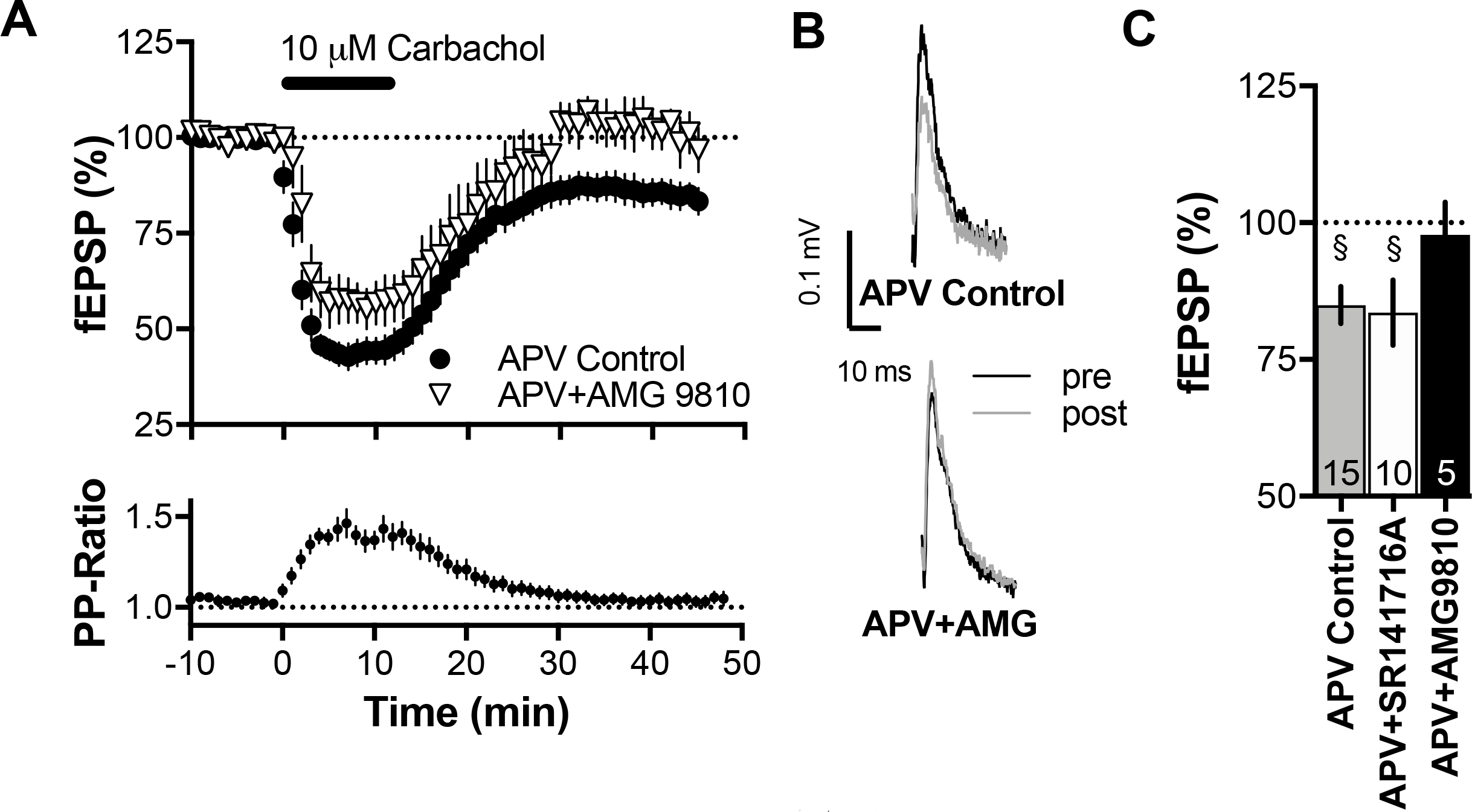
Postsynaptic TRPV1 mediate low Carbachol LTD. A) Averaged field recordings of AMPAR responses showing that preincubation with the TRPV1R antagonist AMG 9810 (10μM) completely prevented the induction of LTD by 10μM Carbachol. B) Example traces of average field response before and after Carbachol application. C) Summary bar graph of the pharmacological experiments characterizing low (10μM) LTD. LTD was blocked by the application of the TRPV1R antagonist AMG9810 (10μM) but not the CB1R antagonist SR141716a (5μM). Error bars represent mean ± s.e.m. § p<0.05, students t-test. The number in each bar indicates the number of experiments

### “High Carbachol” LTD depends on endocannabinoid retrograde signaling and CB1R

We next examined whether CB1R and/or TRPV1 were responsible for high Carbachol/M_1_ mAChR LTD. The CB1R antagonist SR14176A (5μM) efficiently blocked high Carbachol LTD (96.44±5.061, n=7, p=0.5082, student’s t-test Figure 4A-C) whereas the TRPV1 receptor antagonist AMG9810 (10μM) did not prevent the expression of LTD (89.38±1.966, n=7, p=0.0017, n=7; student’s t-test Figure 4A-C). We conclude that high Carbachol LTD requires CB1R, not TRPV1. In the extended amygdala and accumbens both mGluR1 and mGluR5 participate to eCB-LTD^41,45^. In striking contrast, neither the mGluR5 specific antagonist MPEP nor the mGluR1 specific antagonist CPCCoEt prevented from high Carbachol LTD (p = 0.0142; n=8, student’s t-test, data not shown).

**Figure 4:**
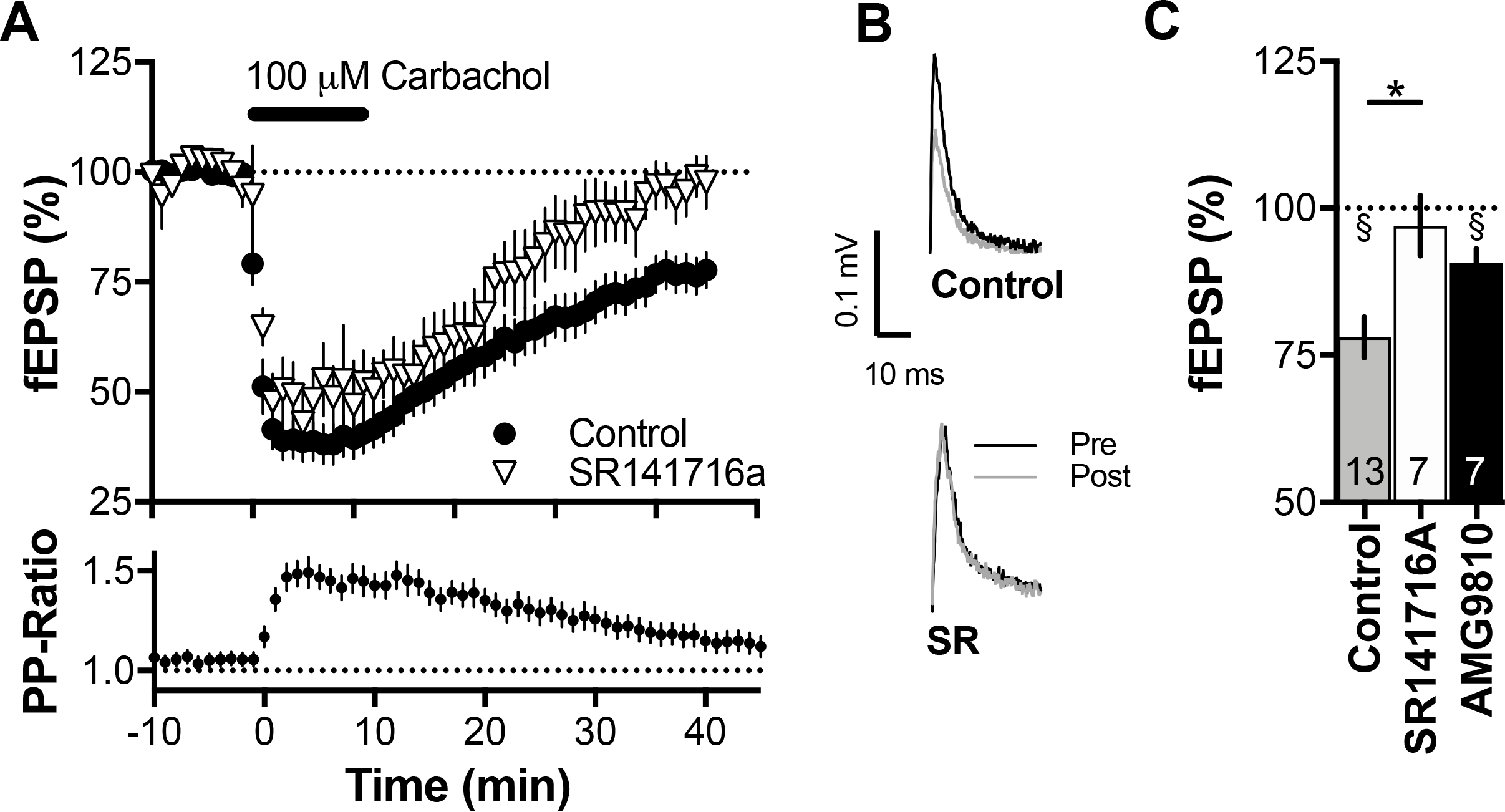
Presynaptic CB1R mediate high Carbachol LTD. A) Averaged field recordings of AMPAR responses. In slices preincubated with the CB1R antagonist SR141716A (5μM), 100μM Carbachol induced STD but not LTD. B) Example traces of average field response before and after Carbachol application. C) Summary bar graph of all pharmacological experiments characterizing the effects of high-Carbachol: LTD was blocked by the application of the CB1R antagonist SR141716a but not by the TRPV1R antagonist AMG9810 (10μM). Error bars represent mean ± s.e.m. * p<0.05, unpaired t-test; § p<0.05, students t-test. The number in each bar indicates the number of experiments.

We verified that high Carbachol LTD had a presynaptic locus of expression as typically expected if CB1R were implicated^44^. Indeed, high Carbachol LTD was paralleled by a significant enhancement of the paired-pulse ratio (p=0.0419, student’s t-test, Figure 4A, lower trace). Together these data suggest that M_1_ mAChR LTD induced by high Carbachol is mediated by eCB acting at presynaptic CB1R.

### CB1R-but not TRPV1-mediated LTD is impaired in fmr1 -/y mice

The postsynaptic mGluR5/eCB signaling complex is impaired at accumbens synapses of *fmr1-/y* mice ^30,31^. M_1_ mAChR and mGluR1/5 are Gq/11-protein coupled receptors with common downstream effectors including eCB^43^. Having established that activation of M_1_ mAChR receptors triggers eCB-mediated LTD via CB1R or TRPV1R; we next tested low and high Carbachol LTD in adult *fmr1-/y* mice. As for Figure 3, the experiments to characterize low Carbachol/TRPV1 R-dependent LTD were performed in the presence of D-AP5 to block NMDAR and unmask LTD. The data show that low Carbachol/TRPV1R-dependent LTD was readily induced in *fmr1-/y* mice (84.21±3.9, n=5, p=0.0155, student’s t-test; Figure 5A) and not different from controls (p=0.743 one-way ANOVA).

**Figure 5:**
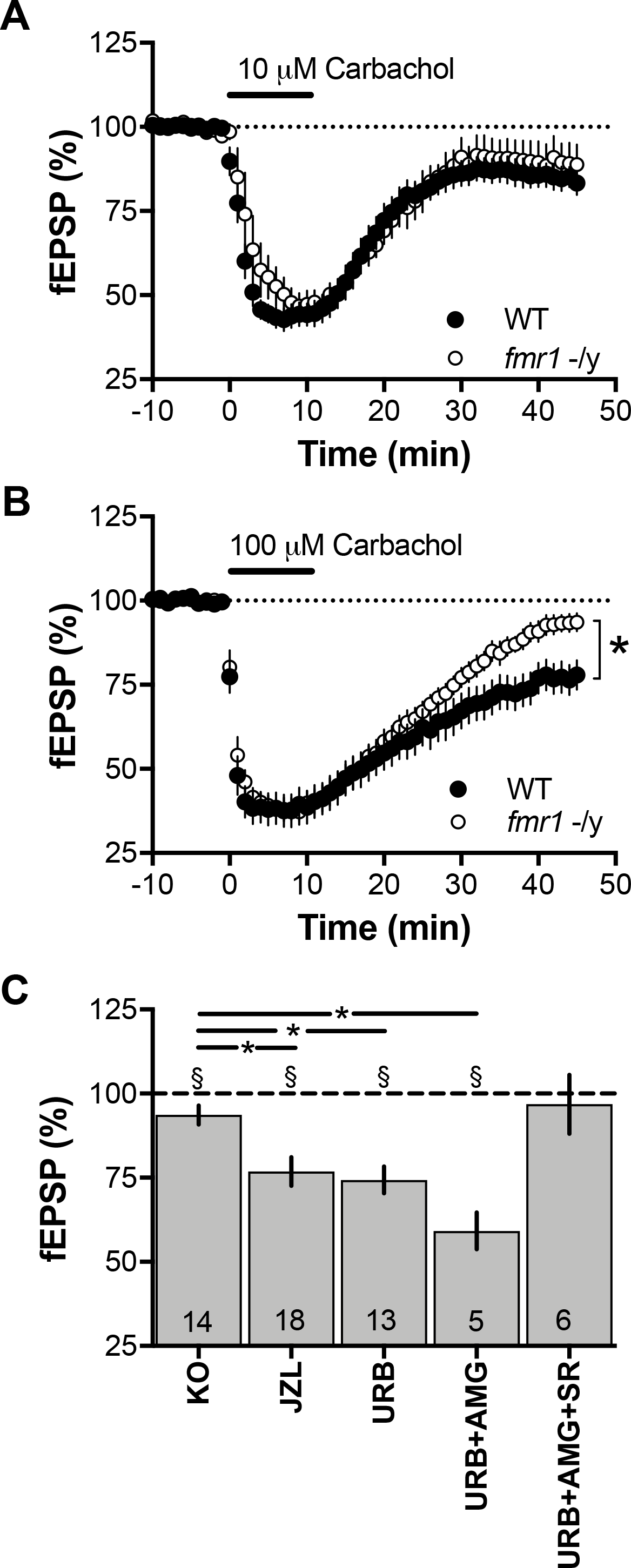
Altered M1R-LTD in *fmr1-/y mice*. A) Summary graph showing that low Carbachol-LTD is intact in *fmr1-/y mice.* B) Summary graph showing that high Carbachol-LTD is abolished in *fmr1-/y mice. C)* Bar graph of pharmacological experiments (mean values of minutes 50-59). The preincubation with URB597 and JZL184 could rescue LTD. LTD rescued by incubation with URB597 could be blocked by SR41716A but not AMG9810. Error bars represent mean ± s.e.m. * p<0.05, unpaired t-test; § p<0.05, students t-test. The number in each bar indicates the number of experiments.

On the contrary, high Carbachol/CB1R-mediated LTD was not abolished (94.55±2.084, n=17, p=0.0187, student’s t-test) but significantly reduced in *fmr1* -/y mice compared to WT littermates (Figure 5B, p=0.0029 unpaired t-test).

In the *fmr1-/y* mouse model, enhancing 2-AG levels by blocking its degradation with the selective monoacylglycerol lipase inhibitor JZL184, normalized synaptic and behavioral impairments^31^. We attempted a similar strategy to rescue deficient high Carbachol LTD in *fmr1-/y.* Indeed, blocking 2-AG degradation with JZL184 restored high Carbachol LTD in *fmr1-/y* mice (Figure 5C, 76.68±4.292, n=15, p<0.001, students t-test).

The two principal signaling eCBs, anandamide and 2-arachidonoylglycerol (2-AG) are thought to have different targets. While anandamide activates CB1R and TRPV1R, 2-AG is thought to mostly engage CB1R ^46–48^ In *fmr1-/y* mice, impaired social interactions and avoidance are improved by URB-597^49,50^, suggesting that elevation of anandamide levels can be beneficial to alleviate from certain Fragile X behavioral symptoms. We tested a similar strategy to relieve the synaptic deficits we had uncovered. In favor of this idea, we found that when accumbens slices from *fmr1-/y* mice were incubated with URB597, high Carbachol treatment induced a LTD indistinguishable from that of wild type littermates (Figure 5C, 74.33±4.041, n=13, <0.001, students t-test). The LTD rescued by incubation with URB597 was not affected by incubation with the TRPV1 antagonist AMG9810 (59.19±5.434, p=0.0017, <0.001, students t-test) but by incubation with the CB1antagonist SR141716A (96.87±8.736 students t-test). These results demonstrate that enhanced anandamide rescues high Carbachol LTD via enhancing CB1R but not TRPV1 signaling.

Neither selective monoacylglycerol lipase inhibitor JZL184 nor the FAAH inhibitor URB-597 had a significant effect on high Carbachol LTD in wildtype littermates (see Supplementary Figure 1).

### Discussion

The principal results of this study are 1/ that at accumbens core synapses, M_1_ mAChR control two forms of endocannabinoid mediated LTD that differ in their signaling pathways and locus of depression and 2/ that lack of FMRP expression selectively impairs CB1R-mediated plasticity.

Low concentration of carbachol engaged TRPV1 receptors and caused a reduction in postsynaptic AMPAR. These results are in agreement of a previous report showing that activation of TRPV1 via the endocannabinoid anandamide induces LTD in the accumbens^45^. Indeed, anandamide has been demonstrated to be an endogenous TRPV1 agonist ^51^

It is important to note that the low Carbachol LTD was unmasked when blocking NMDAR. Several mechanisms could explain the mAChR-mediated synaptic potentiation of NMDAR in accumbens MSN. First, Calabresi et al. have demonstrated that elevation of endogenous acetylcholine increases the conductance of NMDAR^5^. Second, M_1_ -dependent inhibition of SK channels boosts synaptic potentials^40^. Although we cannot unequivocally determine which of these mechanisms mediates NMDAR potentiation, we clearly demonstrate that blocking this potentiation unmasks “low Carbachol” LTD.

The high Carbachol LTD required CB1R-mediated presynaptic inhibition of glutamate release. In the nucleus accumbens core eCB-LTD implicates postsynaptic mGluR5, the production of 2-AG that retrogradely activates presynaptic CB1R^43,44^. Both mGluR5 and M_1_ mAChR are Gq-coupled receptors, that engage similar downstream plasticity mechanisms^52^. M_1_ mAChR regulate inhibitory and excitatory synapses via 2-AG and CB1R^53–57^. Thus, the current data add to the growing list of central synapses where 2-AG is the principal mediator of eCB mediated GPCR synaptic plasticity. Whether low and high Carbachol LTD are induced simultaneously in response to strong M_1_ mAChR stimulation or whether they exclude each other is not completely clear: the slight decrease in LTD after application of a TRPV1 antagonist, which would demonstrate a summation of plasticity did not reach statistical significance (see Figure 4C). The two forms of LTD could engage different signaling pathways by recruiting anandamide for postsynaptic LTD and 2-AG for presynaptic LTD. Indeed, that both eCBs are engaged in M_1_-LTD is supported by the present observation that LTD in *fmr1-/y* mice is rescued by blocking the degradation of either anandamide or 2-AG. How the activation of M_1_ can lead to the engagement of two different endocannabinoid signaling pathways remains to be determined.

Although the production of both endocannabinoids has been shown to depend on GPCR activation, this production can also be state dependent, e.g. depend on activation of voltage gated calcium channels^58^ which could bias the production of one eCB over the other depending on the degree of activation. The two forms of plasticity could also be expressed separately in the two subtypes of medium spiny neurons (i.e. D1R- or D2R-expressing). Although subtype specific synaptic plasticity mechanisms have been reported with various induction protocols ^59,60^ the animal models used have been questioned ^61–64^ The unimodal distribution of LTD observed in patch clamp experiments does not support the idea that CB1R and TRPV1R are expressed in different MSN subtypes (Supplementary Figure 2).

In *fmr1-/y* mice only high CB1R-mediated LTD was ablated, TRPV1R-mediated LTD was normal. Inhibition of either 2-AG or anandamide degradation restored CB1R-LTD. Our results are compatible with recent reports showing that blocking the FAAH inhibitor with URB-597 improves performance in the passive avoidance test and social impairments in *fmr1-/y* mice ^49,50^.

The complex regulation of synaptic plasticity in the accumbens by M_1_ mAChR supports the idea that the cholinergic system is a substrate of arousal and emotional deficits observed in Fragile X.

## Methods

### Animals

Animals were treated in compliance with the European Communities Council Directive (86/609/EEC) and the United States National Institutes of Health Guide for the Care and Use of Laboratory Animals. All animals were housed with 12 h light/dark cycles and access to food and water ad libitum.

### Slice Preparation

Adult male *fmr1-/y* mice on a C57Bl6/J genetic background aged between 60 and 95 postnatal days were used, with wild-type littermates and C57Bl6/J mice purchased from Janvier Labs France used as control group^31^. They were anesthetized with isoflurane and decapitated according to institutional regulations. The brain was sliced (300 μm) in the coronal plane with a vibratome (Integraslice, Campden Instruments, Loughborough, UK) in a sucrose-based solution at 4°C (in mM: 87 NaCl, 75 sucrose, 25 glucose, 2.5 KCl, 4 MgCl_2_, 0.5 CaCl_2_, 23 NaHCO_3_ and 1.25 NaH_2_PO_4_). Immediately after cutting, slices were stored for one hour at 32°C in a low calcium artificial cerebrospinal fluid (low Ca^2+^ ACSF) that contained (in mM): 130 NaCl, 11 Glucose, 2.5 KCl, 2.4 MgCl_2_, 1.2 CaCl_2_, 23 NaHCO_3_, 1.2 NaH_2_PO_4_, and was equilibrated with 95% O_2_/5% CO_2_. Slices were maintained at room temperature until the time of recording.

### Electrophysiology

Field potential recordings were made in coronal slices containing the accumbens core as previously described^30,44^. Recordings were made in the medial ventral accumbens core close to the anterior commissure^30, 44^.

For recording, slices were placed in the recording chamber and superfused (1.5 - 2 ml/min) with ACSF (same as low Ca^2+^ ACSF with the following exception: 2.4 mM CaCl_2_ and 1.2 mM MgCl_2_). All experiments were done at 25°C. Picrotoxin (100 μM) was added to the superfusion medium to block gamma-aminobutyric acid type A (GABA-A) receptors. All drugs were added at the final concentration to the superfusion medium (see Table 1).

**Table 1.**
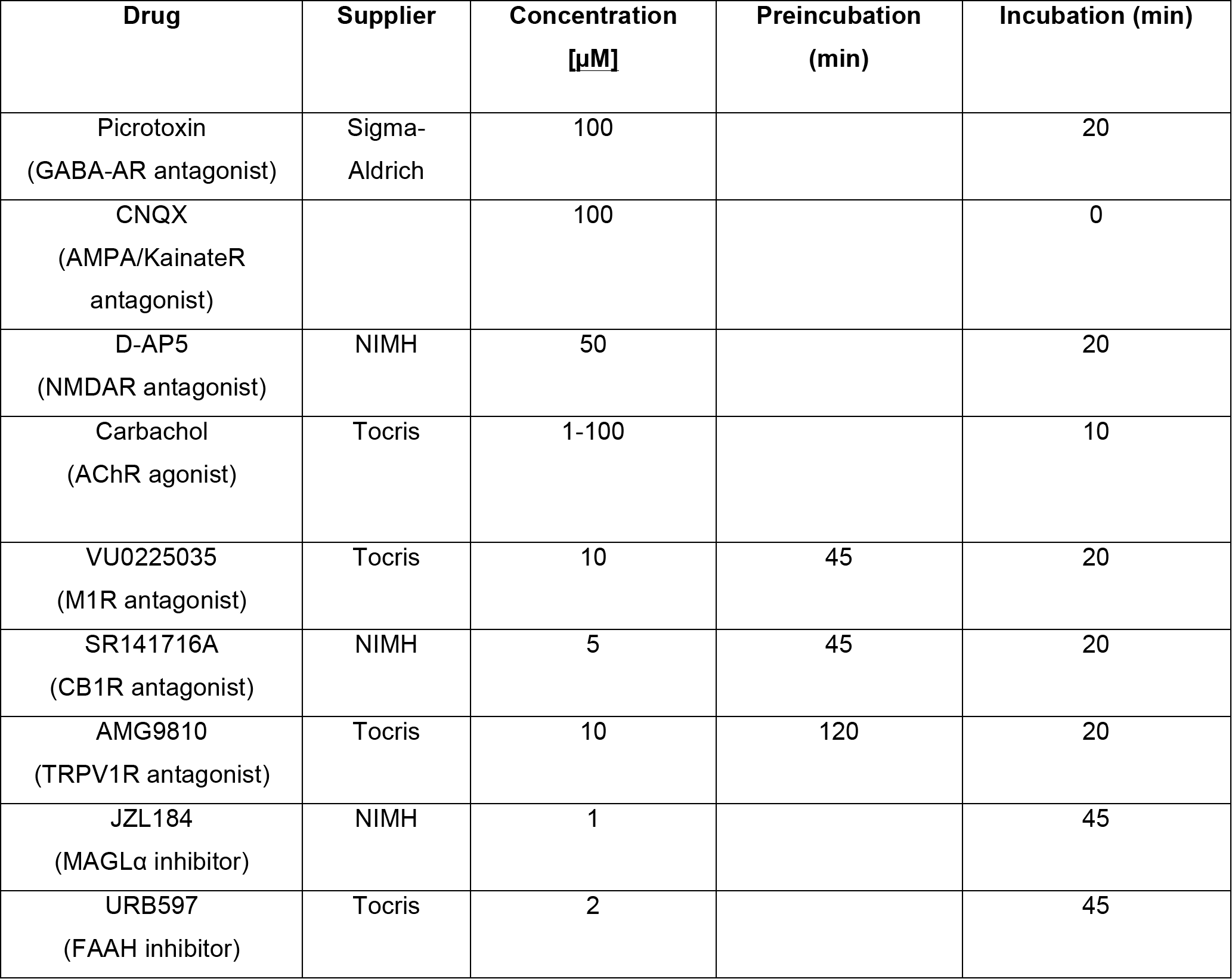
Drugs suppliers, final concentrations and incubation times.

For field excitatory postsynaptic potential (fEPSP), the recording pipette was filled with ACSF and afferents were stimulated with a glass electrode filled with ACSF and placed ~200μm in the dorsal-medial direction of the recording pipette. The stimulus intensity was adjusted around 80% of maximal intensity after performing an input-output curve (baseline fEPSP amplitudes ranged between 0.15 mV and 0.4 mV). Stimulation frequency was set at 0.1 Hz.

Recordings were performed with an Axopatch-200B amplifier (Axon Instrument, Molecular Device, Sunnyvale, USA). Data were lowpass filtered at 2kHz, digitized (10kHz, DigiData 1440A, Axon Instrument, Molecular Device, Sunnyvale, USA), collected using Clampex 10.2 and analyzed using Clampfit 10.2 (Axon Instrument, Molecular Device, Sunnyvale, USA). Both fEPSPs’ area and amplitude were analyzed.

### Drugs

All drugs were added at final concentration to the superfusion medium (see Table 1 for details).

### Data acquisition and analysis

The magnitude of plasticity was calculated 40-44 minutes after the plasticity protocols as percentage of baseline responses. Statistical analysis of data was performed with GraphPad Prism (GraphPad Software Inc., La Jolla, CA). All values are given as mean ± standard error. N indicates the number of experiments. At least 3-12 animals were used for each single experimental condition. The Shapiro-Wilk test confirmed the normal distribution of data sets. Therefore, depending on the experimental condition, statistical differences were assessed using t-tests and one-way ANOVAs + Fisher LSD Post hoc tests. A confidence level of P < 0.05 was considered significant.

## Author contributions

D.N. and O.J.M. designed research; D.N. and O.L. performed research; D.N. and O.L. analyzed data; D.N. and O.J.M. wrote the paper. The authors declare no conflict of interest.

## Acknowledgments

The FRAXA Foundation (O.J.M. and D.N.), a NARSAD 2010 Independent Investigator Grant given by the Brain & Behavior Research Foundation (O.J.M.), Agence Nationale de la Recherche (ANR CortexCell and Cyfip-Aut; O.M. and O.L.) and INSERM (OJ.M.) supported this work. The authors acknowledge FRAXA research foundation (Dr. D. Nelson, Baylor College of Medicine) for providing the Fmr1 KO2 mice. We thank members from the O. Manzoni and P. Chavis laboratories for discussions and the National Institute of Mental Health’s Chemical Synthesis and Drug Supply Program (Rockville, MD, USA).

---

**Supplementary Figure 1.**
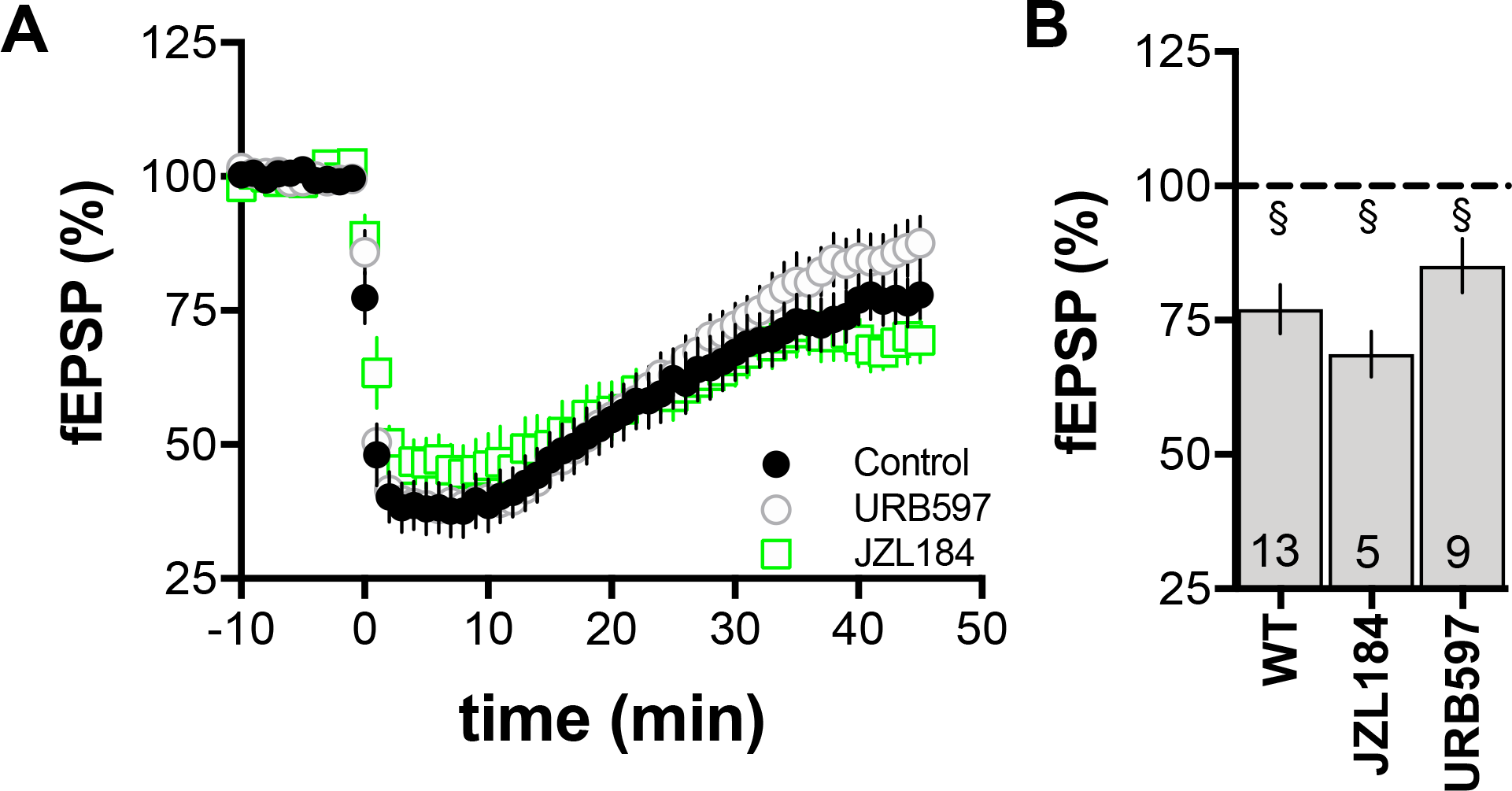
Effect of Blockade of 2-AG and Anandamide degradation on high Carbachol LTD in slices from Wild type littermates. A,B) Summary graph showing that high Carbachol-LTD is inot changed in slices incubated with JZL184 or URB597 B) Summary graph showing that high Carbachol-LTD is abolished in *fmr1-/y mice*. Error bars represent mean ± s.e.m. * § p<0.05, students t-test. The number in each bar indicates the number of experiments.

**Supplementary Figure 1.**
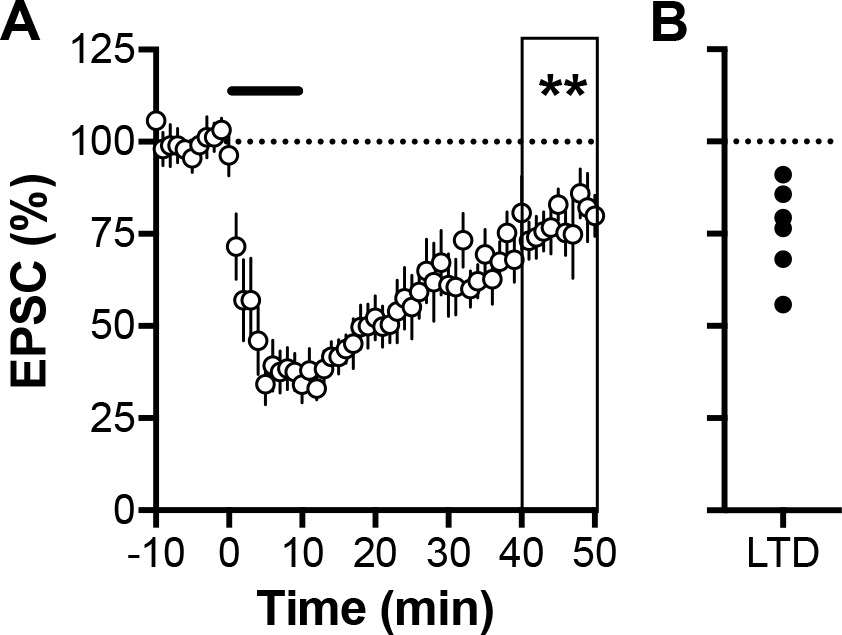
Whole Cell patch clamp recordings of accumbens MSNs. **A)** Averaged patch clamp recordings of AMPAR responses. 100μM Carbachol induced strong STD and LTD. B) The average LTD values of this dataset indicate that all cells recorded expressed LTD. For whole cell patch-clamp experiments, neurons were visualized using an upright microscope (Olympus BX-51W) with infrared illumination. The intracellular solution contained in mM: 145 K^+^ gluconate, 3 NaCl, 1 MgCl_2_, 1 EGTA, 0.3 CaCl_2_, 2 Na_2_^+^ATP, and 0.3 Na^+^ GTP, 0.2 cAMP, buffered with 10 HEPES. The pH was adjusted to 7.2 and osmolarity to 290-300 mOsm. Electrode resistance was 4-6 MOhms. A −2 mV hyperpolarizing pulse was applied before each evoked EPSC in order to evaluate the access resistance and those experiments in which this parameter changed >25% were rejected. Access resistance compensation was not used and acceptable access resistance was <30 MOhms. The potential reference of the amplifier was adjusted to zero prior to breaking into the cell. Cells were held at −70mV.

